# Microcystin-Driven Control of the Carbon-Concentrating Mechanism Shapes CO_2_ Fixation Dynamics in *Microcystis aeruginosa* PCC 7806

**DOI:** 10.64898/2026.07.10.737655

**Authors:** Arthur Guljamow, Stefan Timm, Vivienne Wimmer, Luca Schulz, Georg Hochberg, Martin Hagemann, Elke Dittmann

## Abstract

Bloom-forming cyanobacteria thrive in highly dynamic light environments, yet the mechanisms enabling rapid acclimation to fluctuating irradiance remain poorly understood. Here, we compared light acclimation in the bloom-forming cyanobacterium *Microcystis aeruginosa* PCC 7806 and the non-bloom-forming model cyanobacterium *Synechocystis* sp. PCC 6803 and investigated the role of the cyanobacterial toxin microcystin (MC) and its *in vivo* binding partner RubisCO in this process. Whereas *Synechocystis* grew faster under sustained high light, *Microcystis* performed better under low light and responded to transient high-light exposure with a remarkably rapid increase in photosynthetic activity and glycogen accumulation. These responses were markedly attenuated in an MC-deficient mutant. Although RubisCO from *Microcystis* exhibited pronounced light-dependent changes in activity, MC had only minor effects on RubisCO catalysis, arguing against a direct role in regulating enzyme function. Instead, extracellular MC elicited a transient transcriptional program characterized by induction of inorganic carbon acquisition systems, including the high-affinity bicarbonate transporter BCT1, consistent with activation of the carbon-concentrating mechanism (CCM) and enhanced carbon fixation *in vivo*. MC further stimulated the expression of photosynthesis-related genes, and altered carboxysome organization, and promoted extracarboxysomal localization of RubisCO. Together, our findings identify MC as a light-responsive signaling molecule that coordinates CCM activity, carbon acquisition, and photosynthetic acclimation, thereby enhancing adaptation of *Microcystis* to fluctuating irradiance and potentially contributing to its ecological success in cyanobacterial blooms.

## Introduction

Cyanobacteria are integral to freshwater ecosystems, supporting primary production and nutrient cycling, yet excessive bloom formation leads to a deterioration in water quality, reduces biodiversity, and impairs fisheries and recreation (*1*). *Microcystis* is an important taxon that forms harmful algal blooms (cyanoHABs), with its negative impacts primarily attributable to the production of microcystins (MCs)—hepatotoxins that threaten drinking water supplies, and human health (*2*). *Microcystis* spp. exhibits pronounced genotypic and phenotypic plasticity, enabling rapid adaptation to changing environmental conditions, including buoyancy regulation, a key trait for bloom formation (*3*). This flexibility allows it to optimize access to light and nutrients within the water column and to persist under highly variable environmental conditions. Genomic plasticity in *Microcystis* spp. notably includes variations in bicarbonate transport genes, diversifying its capacity for inorganic carbon acquisition under changing conditions (*4*). It also encompasses diverse biosynthetic gene clusters for a specialized metabolism, including those responsible for MC production (*5*). Single-colony sequencing identified at least 18 distinct genotypes within the phylum *Microcystis*, with MC production unevenly distributed among them (*6*). Because bloom toxicity largely depends on the proportion of toxic genotypes, differences in the adaptive strategies of toxic and non-toxic *Microcystis* strains have been used to predict bloom toxicity under climate change conditions and to inform management strategies (*7*). Although MC is toxic to potential grazers, its primary role as a defense against eukaryotes is increasingly questioned, partly because its early evolutionary origin suggests additional physiological or ecological functions (*8*).

Genomic variability and the marked differences observed among various *Microcystis* strains in their adaptation to different inorganic carbon conditions are particularly noteworthy given growing evidence linking MC to the regulation of photosynthetic CO_2_ fixation. For example, the MC-producing wild-type strain *M. aeruginosa* PCC 7806 and the MC-deficient mutant Δ*mcyB* differ markedly in their ability to reduce dissolved CO₂ at low concentrations (*9*) and exhibit transcriptomic, proteomic, and metabolomic differences related to the Calvin–Benson–Bassham (CBB) cycle and bicarbonate uptake transport (*10–12*). These differences are most pronounced during shifts from low-light to high-light conditions, with the MC-deficient mutants displaying a distinct high-light phenotype (*12*). While the mechanistic role of MC under high-light conditions remains unresolved, two MC-related phenomena have been observed under high-light conditions: increased binding of MC to proteins, including the key enzyme for CO_2_ fixation, RubisCO (*12*), and the secretion of small amounts of MC (*13*). Both phenomena exhibit pronounced diurnal fluctuations (*14*) which were shown to be correlated with a non-canonical extracarboxysomal localization of RubisCO beneath the cytoplasmic membrane in the MC-producing wild type and also involve a recurrent turnover of RubisCO (*15*).

Here, we investigated the role of microcystin and its effects on RubisCO both *in vitro* and *in vivo*, while also incorporating the non-bloom-forming model cyanobacterium *Synechocystis* sp. PCC6803 into our analyses. Our findings indicate that MC does not directly affect RubisCO activity *in vitro*; however, extracellular MC appears to enhance CO_2_ fixation *in vivo*, at least in part through the induction of genes encoding bicarbonate uptake transporters and by influencing RubisCO localization. Collectively, our results highlight the complex interplay between MC, the carbon concentrating mechanism (CCM), and RubisCO, and provide new mechanistic insights into the non-canonical high-light acclimation strategies of the notorious bloom-forming cyanobacterium *Microcystis*.

## Results

### Rapid High-Light Acclimation Distinguishes Microcystis from Synechocystis and Depends on Microcystin

To assess the effect of MC on low– and high-light acclimation of the bloom-forming cyanobacterium Microcystis, we monitored growth of the laboratory strain Microcystis aeruginosa PCC7806 and its MC-free ΔmcyB mutant. As a control we included the non-bloom-forming model cyanobacterium Synechocystis sp. PCC6803 and its fully segregated rbcLXS replacement mutant variant (RMA) that expresses the RubisCO coding sequence from Microcystis aeruginosa PCC7806 from a chromosomal integration at the native rbc locus (Figs. 1A and S1). Under low-light conditions, both Microcystis strains grew significantly faster than either Synechocystis strain (Figs. 1B and C). However, MC had no detectable effect on the strong low-light performance of Microcystis. Growth of the RMA strain largely mirrored that of the wild-type Synechocystis strain but was accompanied by the development of a chlorotic phenotype after 10 days (Fig. 1B). In contrast, under high light, both Synechocystis strains markedly outperformed Microcystis with the RMA strain entering into stationary phase earlier than the wild type. Consistent with previous reports (12), growth of the MC-deficient Microcystis ΔmcyB mutant was significantly impaired relative to the wild type under high light.

**Fig. 1:**
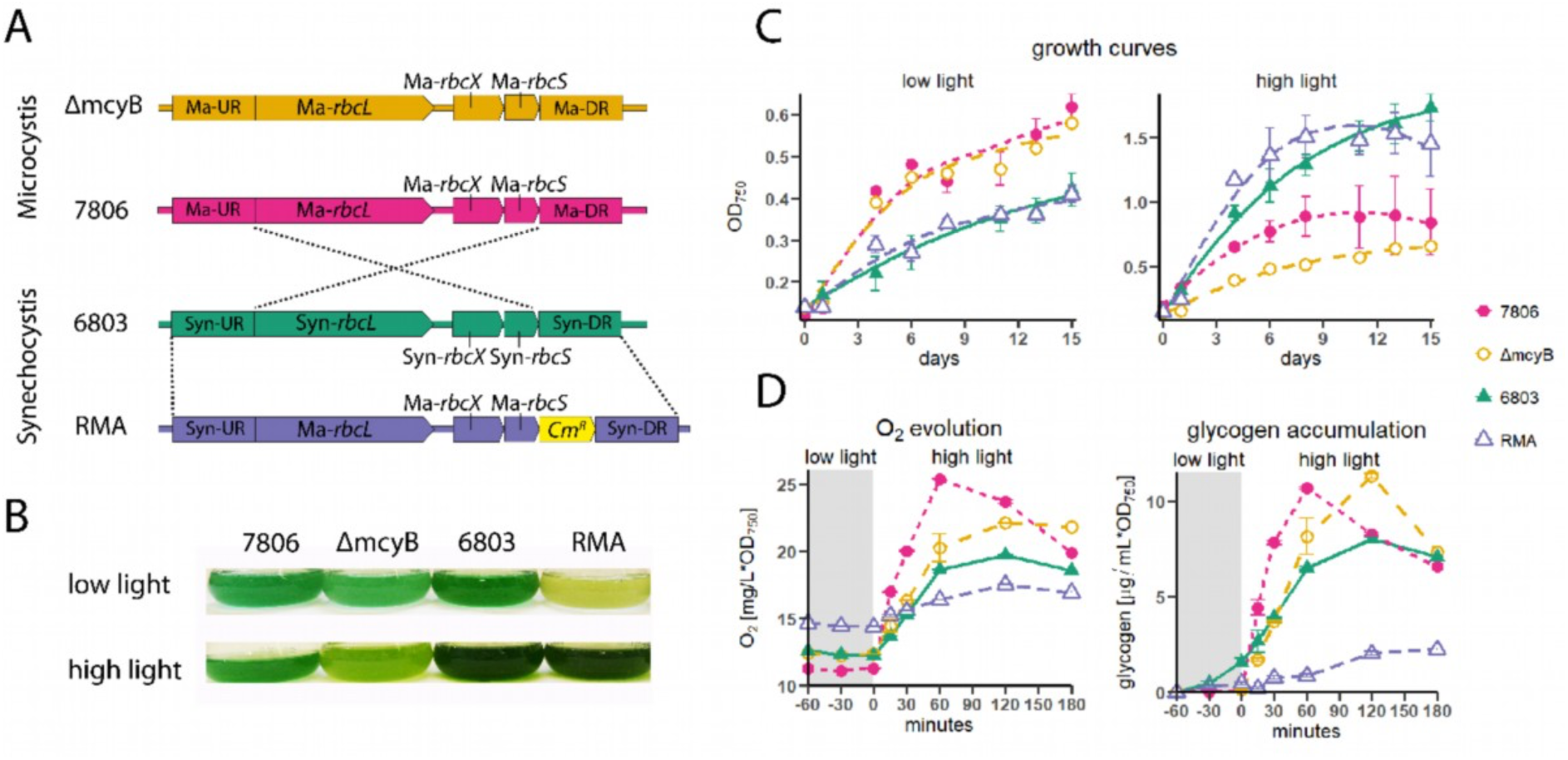
Light-dependent acclimation of cyanobacteria harboring different RubisCO forms. (**A**) Schematic representation of the *rbcLXS* genomic locus encoding RubisCO in the four genotypes used in this study. Strain codes: “7806”, *Microcystis aeruginosa* wild type; “ΔmcyB”, microcystin-deficient mutant; “6803”, *Synechocystis* sp. wild type; and “RMA”, a *Synechocystis* RubisCO replacement mutant expressing the *Microcystis rbcLXS* variant. (**B**) Photographs of liquid cultures of the four cyanobacterial strains grown for 10 days under low light (15 µmol photons m⁻² s⁻¹) or high light (150 µmol photons m⁻² s⁻¹). (**C**): Growth curves of liquid cultures under continuous low-light or high-light conditions based on optical density measurements. (**D**) Quantification of oxygen evolution and glycogen accumulation during a short-term shift from low light (15 µmol photons m⁻² s⁻¹) to high light (250 µmol photons m⁻² s⁻¹) in all four strains.

In addition to the long-term growth analyses under continuous illumination, we examined the short-term response of all four genotypes to a shift from low to high light. To this end, we monitored oxygen evolution as a proxy for photosynthetic activity and glycogen accumulation as indicator for carbon assimilation (Fig. 1D). Following transfer to high light, *Microcystis* very rapidly increased its photosynthetic activity and concomitantly exhibited fast and strong glycogen accumulation. Both responses were delayed and attenuated in the MC-deficient mutant. In contrast, *Synechocystis* did not display a similarly dynamic response but instead exhibited a more gradual and stable, albeit weaker, acclimation to high light.

Notably, the rapid response of *Microcystis* was transient. After only one hour of high-light exposure, both oxygen evolution and glycogen accumulation declined sharply. This decline was likewise delayed and attenuated in the MC-deficient mutant. In both assays, the RMA strain showed severe impairment compared to the other strains. This indicates that despite the high sequence conservation of RubisCO between *Microcystis* and *Synechocystis* (97.2% and 84.4% similarity for RbcL and RbcS, respectively), the *Microcystis* variant cannot serve as a full functional replacement of the *Synechocystis* form. Taken together these findings show that MC plays a role both in long-term and short-term high-light acclimation of the bloom-forming cyanobacterium *Microcystis*. The exceptionally rapid short-term response to high light is particularly noteworthy, as *Microcystis* clearly outperforms *Synechocystis* during the initial high light acclimation phase.

### Microcystin does not enhance RubisCO activity in vitro, but stimulates CO_2_ fixation in vivo

To determine whether the pronounced differences in light acclimation could be attributed to intrinsic differences in RubisCO catalysis, we purified the enzyme from all four genotypes (Fig. 2A) and characterized its biochemical properties. All four RubisCO forms were recovered in their canonical L_8_S_8_ holoenzyme conformation (Fig. 2B). Moreover, the three enzymes carrying the *Microcystis* amino acid sequence exhibited similarly high CO₂/O₂ specificity factors, among the highest reported for cyanobacterial RubisCOs (Fig. 2C). Importantly, these specificity factors were independent of whether RubisCO was isolated from MC-producing cells (wild-type *Microcystis*) or from MC-free backgrounds (Δ*mcyB* and RMA), indicating that MC does not directly alter the intrinsic catalytic properties of the enzyme. To test this more directly, we measured RubisCO carboxylase activity by quantifying 3-phosphoglycerate (3PGA) production in enzyme assays performed in the presence or absence of purified MC. In parallel, MC-binding to the two *Microcystis* RubisCO variants was confirmed by immunoblot analysis (Fig. S2). We found an *in vitro* inhibiting effect of MC on wild-type *Microcystis* RubisCO but only negligible effects on all other RubisCO preparations (Fig. 2D). As these RubisCOs were isolated from high-light grown cultures, we next compared 3PGA levels produced by RubisCO isolated from cells grown in parallel under either low light or high light. Whereas *Synechocystis* RubisCO showed little variation in activity between the two light regimes, *Microcystis* RubisCO displayed pronounced light-dependent plasticity, particularly in the MC-deficient Δ*mcyB* mutant. RubisCO isolated from low-light-grown Δ*mcyB* cells exhibited substantially higher activity than the enzyme isolated from cells grown under high light. Notably, RubisCO purified from the RMA strain did not reproduce this light-dependent activity pattern under either condition. This observation is particularly striking because the RubisCO enzymes from Δ*mcyB* and RMA are identical in amino acid sequence and were both isolated from cells that had never been exposed to MC. Nevertheless, they exhibited marked differences in carboxylase activity, especially when purified from low-light-grown cultures. These findings indicate that factors beyond MC itself contribute to the observed differences in RubisCO activity.

**Fig. 2:**
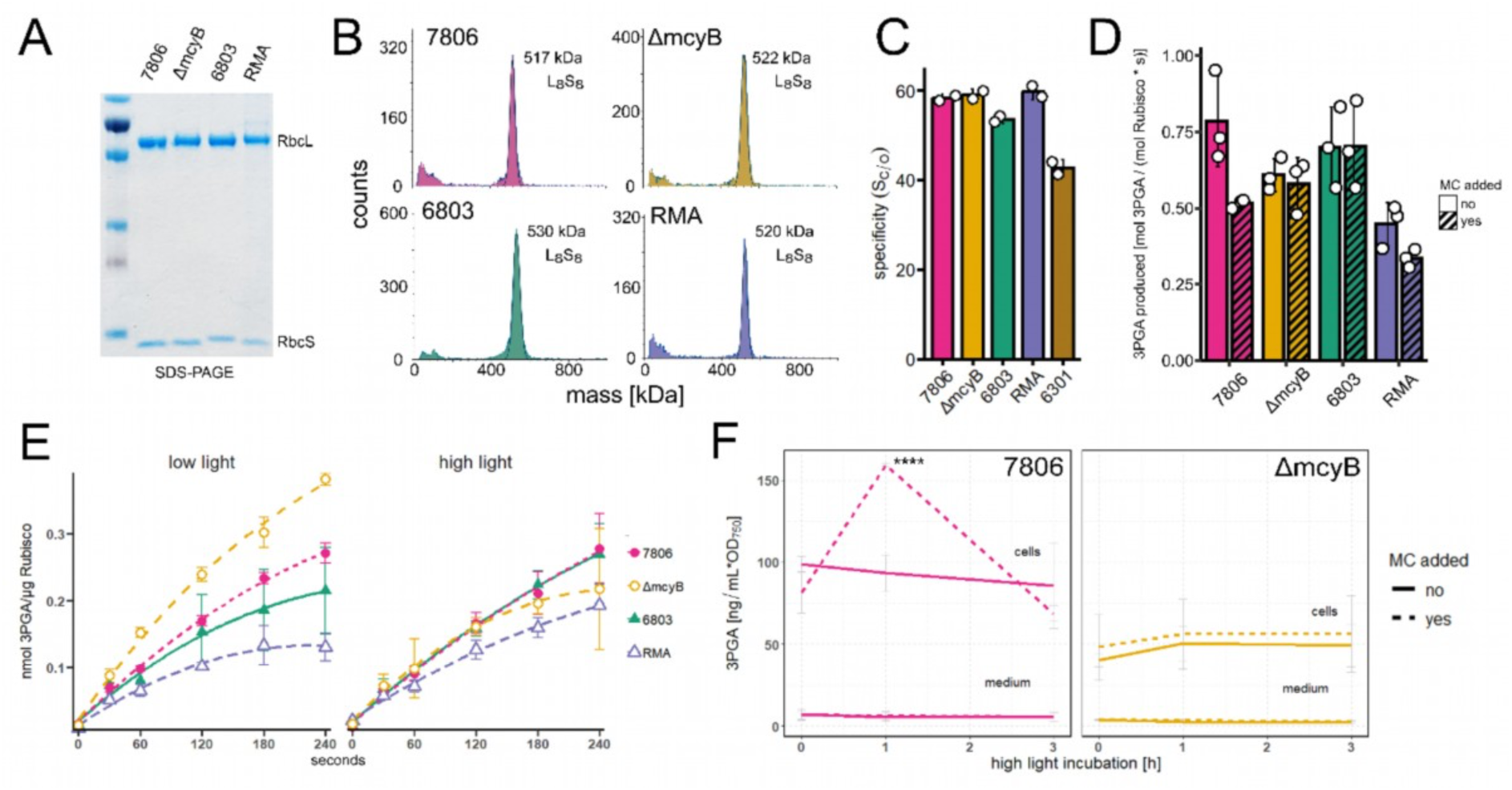
Effect of microcystin on RubisCO activity *in vitro* and *in vivo*. (**A**) Coomassie-stained SDS-PAGE gel of purified RubisCO complexes from the four cyanobacterial genotypes. The large (RbcL) and small (RbcS) subunits are indicated. (**B**) Mass photometry analysis of native RubisCO complexes, showing L₈S₈ holoenzyme formation in all tested variants. (**C**) Substrate specificity factor (S_C/O; CO₂ vs. O₂) of RubisCO enzymes isolated from different cyanobacteria. *Synechococcus elongatus* PCC 6301 RubisCO served as a reference. (**D**) Effect of *in vitro* addition of microcystin on RubisCO carboxylase activity. (**E**) RubisCO activity assays using purified enzymes from cyanobacteria grown under low-light (left) and high-light (right) conditions. (**F**) Effect of externally added microcystin on intracellular (“cells”) and extracellular (“medium”) 3-phosphoglycerate (3PGA) levels in high-light-shifted *Microcystis* cultures. Asterisks indicate ANOVA significance (p < 0.001).

We next investigated whether extracellular MC could influence CO_2_ fixation *in vivo*. To this end, MC was added to wild-type *Microcystis* cultures at a concentration of 100 ng mL⁻¹, corresponding to extracellular levels detected during diurnal experiments (*14*). We then monitored the accumulation of the RubisCO product 3PGA following a shift from low to high light irradiation. Remarkably, MC addition triggered a rapid but transient increase in intracellular 3PGA levels, peaking approximately one hour after treatment (Fig. 2F). This response largely corresponded to the transient peak in photosynthetic oxygen evolution and glycogen accumulation previously observed in wild-type cells during low– to high-light shifts (Fig. 1D).

Together, these results demonstrate that MC does not directly modulate RubisCO catalysis *in vitro*. Instead, its stimulatory effect on CO_2_ fixation appears to occur *in vivo*, potentially through extracellular signaling mechanisms that regulate RubisCO activity or, more broadly, carbon metabolic processes in response to changing light conditions.

### Extracellular microcystin coordinates carbon acquisition and photosynthetic responses during high-light acclimation

To further elucidate the role of MC in the high-light acclimation of *Microcystis*, we sequenced the transcriptomes of the wild type and Δ*mcyB* mutant in the presence or absence of externally added MC under either low-light conditions or following a shift from low to high irradiance. To capture the pronounced dynamics observed after the high-light transition (Figs. 1D and 2F), samples were collected 1 and 3 h after the shift. Under both low– and high-light conditions, the wild type and Δ*mcyB* mutant exhibited distinct transcriptional profiles, as reflected by their clear separation in principal coordinate analyses (Fig. 3A).

**Fig. 3:**
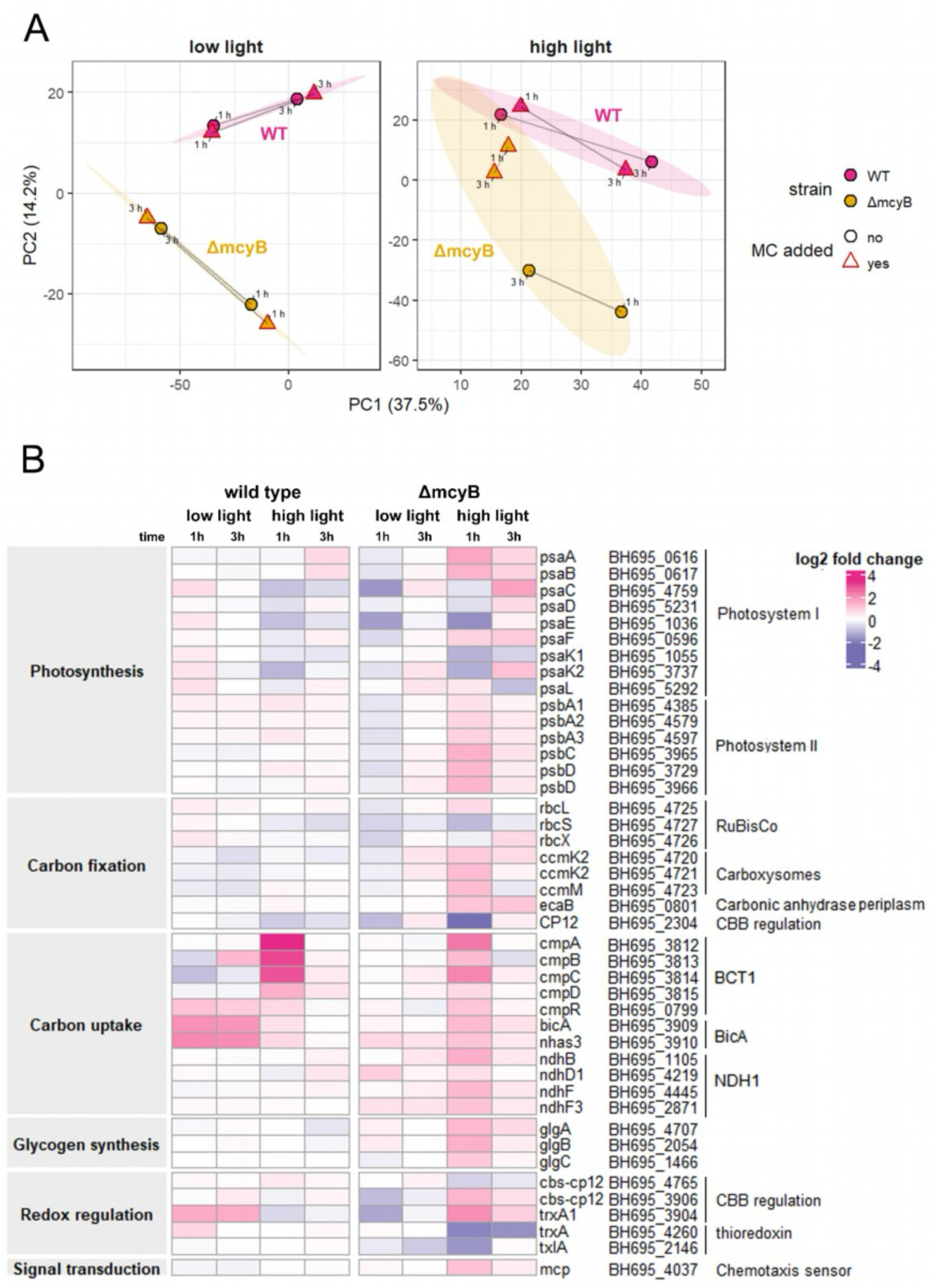
Transcriptional response of *Microcystis* to externally added microcystin. (**A**) Principal coordinate analysis of transcriptome data from the microcystin addition experiment based on TPM (transcripts per million) values. *Microcystis aeruginosa* PCC 7806 wild type and its Δ*mcyB* mutant were grown under low-light and high-light conditions for 1 and 3 h, with or without microcystin supplementation. Statistical ellipses represent a 0.75 confidence level; lines connect time points (1 and 3 h). MC addition is indicated by red triangles. “WT”: *M. aeruginosa* PCC 7806 wild type; “ΔmcyB”: microcystin-deficient mutant. (**B**) Heat map of differential gene expression showing log₂ fold changes in transcript levels between MC-treated and untreated cells under low-light and high-light conditions over 1 and 3 h. Functional COG categories are shown in shaded boxes on the left. Gene names, locus tags, and functional annotations are listed on the right.

Addition of MC had virtually no effect under low-light. In contrast, under high light it induced extensive transcriptional reprogramming in the Δ*mcyB* mutant. Notably, MC-treated mutant samples shifted toward the wild-type high-light transcriptome, suggesting that extracellular MC can at least partially restore the wild-type transcriptional response to high light (Fig. 3A). Analysis of MC-responsive genes across COG functional categories confirmed that the strongest transcriptional effects occurred in the mutant under high-light exposure, consistent with a broad cellular response to MC. This response was already markedly attenuated after 3 h, highlighting the transient nature of MC-mediated regulation (Fig. S3, dataset S1). Among the four most affected categories, Mobilome (X), Defense Mechanisms (V), and General Function Prediction Only (R) were predominantly downregulated following MC addition, while the category Energy Production and Conversion showed a pronounced positive response, particularly after 1h of high-light exposure (Fig. S3, dataset S1).

Genes encoding RubisCO itself were largely unaffected, although *rbcS* was slightly downregulated. Most strikingly, extracellular MC strongly affected carbon acquisition and CCM-related genes. After 1 h of high-light exposure, MC addition induced the major inorganic carbon uptake systems (*bct1*, *bicA*, and to a lesser extent *ndhI*), carboxysome-associated *ccm* genes, the periplasmic carbonic anhydrase gene *ecaB*, and regulatory genes of the CBB cycle (Fig. 3B). A notable exception was *cp12*, which was strongly downregulated. Because CP12 represses key CBB-cycle enzymes, its downregulation is consistent with enhanced carbon fixation and activation of the carbon-concentrating machinery (*16*). MC also stimulated the expression of many photosynthesis-related genes in the mutant under high-light conditions, including the photosystem I reaction center genes *psaAB* and several photosystem II components (Figs. 3B and S4). At the same time, several accessory photosystem I genes, particularly *psaC*, *psaE*, and *psaK*, were repressed. Consistent with the upregulation of photosynthesis and carbon uptake, genes involved in glycogen biosynthesis were also induced under high light in mutant cells (Figs. 3B and S4). In the wild type, by contrast, only a small number of genes responded to externally added MC. These included a strong upregulation of the high affinity bicarbonate transporter BCT1 and a marked downregulation of some photosystem 1 genes in response to MC addition after a high light shift (Fig. 3B). The low affinity, constitutive bicarbonate transporter BicA was also MC-responsive in the wild type, albeit only under low-light conditions, where its expression increased. (Fig. 3B). The generally weaker effect of MC on the wild-type strain likely reflects its native exposure to external MC rendering it less responsive to additional MC (Fig. 3A).

Another group of MC responsive genes comprised components of the cellular redox regulatory network, particularly thioredoxins. Two trxA genes were repressed by MC, whereas a third trxA homolog was induced. Interestingly, this induced trxA gene is associated with a CBS-CP12 gene that also responds to MC and was recently characterized as a regulator of the CBB cycle (*17*). We also carefully evaluated the COG category Signal Transduction for candidates involved in MC-dependent signaling. Apart from the carbon-responsive CmpR regulator (Fig. 3B), which controls the BCT1 system, no obvious regulators capable of explaining the observed global transcriptional reprogramming were identified (dataset S1). However, there was a putative methyl-accepting chemotaxis regulator, mcp, which was upregulated in the mutant under high-light conditions following MC addition and could represent a candidate for an MC receptor (Fig. 3B).

Comparison of the high-light response in *Microcystis* in the presence and absence of MC suggests that several hallmark acclimation processes (*18*) are accelerated or amplified by MC. These include earlier induction of the bicarbonate uptake systems BCT1 and BicA, enhanced expression of photosystem II genes such as *psbA*, and modulation of the photosystem I/photosystem II balance through differential regulation of PSI components (Fig. S4). Together, these findings suggest that extracellular MC triggers a rapid but transient stimulation of photosynthesis and carbon acquisition. Although this response may reduce the extent of PSI downregulation that normally limits photodamage under high light, repression of specific PSI subunits such as PsaC and PsaE may still contribute to protection against excess light stress. Several aspects of the MC response also differ from canonical high-light acclimation responses described in other cyanobacteria. Most notably, high-light-induced expression of *cp12*—previously reported in *S. elongatus* PCC 7942 and likewise observed in the *Microcystis* strains examined here—was suppressed by MC (*19*). In *Microcystis*, however, this response may be functionally compensated by the concurrent upregulation of a CBS-CP12-encoding gene, whose product appears to be at least partially redundant with CP12 in regulating the CBB cycle. Overall, the transcriptomic data support a model in which extracellular MC acts as a transient signal that promotes carbon acquisition, photosynthetic activity, and high-light acclimation, thereby facilitating the rapid adaptation of *Microcystis* to fluctuating light conditions.

### External microcystin affects RubisCO localization and carboxysome organization in vivo

To determine whether externally added MC influences key CCM components beyond gene expression, we analyzed the localization of RbcL (the large subunit of RubisCO, Fig. 4A-C) and CcmK (the major structural component of the carboxysome, Fig. 4D-F) by immunofluorescence microscopy. Specifically, we asked whether exogenous MC could restore the differences in RubisCO localization previously observed between the wild type and the MC-deficient mutant (*15*). RubisCO localization was highly plastic, particularly in the MC-producing wild type. We classified the observed patterns into four localization states: (i) carboxysomal, (ii) diffuse extracarboxysomal, (iii) mixed carboxysomal/extracarboxysomal, and (iv) peripheral (Fig. 4A), with peripheral localization typically occurring together with diffuse or mixed patterns. The MC-deficient mutant showed predominantly carboxysomal RubisCO under both light conditions (Fig. 4B). In contrast, the wild type displayed substantially more mixed carboxysomal/extracarboxysomal and peripheral localization (Fig. 4A). While the overall localization pattern in the wild type was similar under low and high light, the extent of peripheral RubisCO localization increased under high light (Fig. 4A). Consistent with the increased extracarboxysomal RubisCO pool, carboxysome shell genes were downregulated in the wild-type strain under high-light conditions (Fig. S4). The addition of MC had only modest effects on RubisCO localization in the wild type, causing a gradual reduction in peripheral localization. In contrast, MC treatment elicited a distinct response in the ΔmcyB mutant under high light. Alongside carboxysomal localization, most cells developed pronounced a diffuse extracarboxysomal RubisCO localization, partially restoring the wild-type phenotype. Notably, however, peripheral RubisCO localization was not reconstituted.

**Fig. 4.**
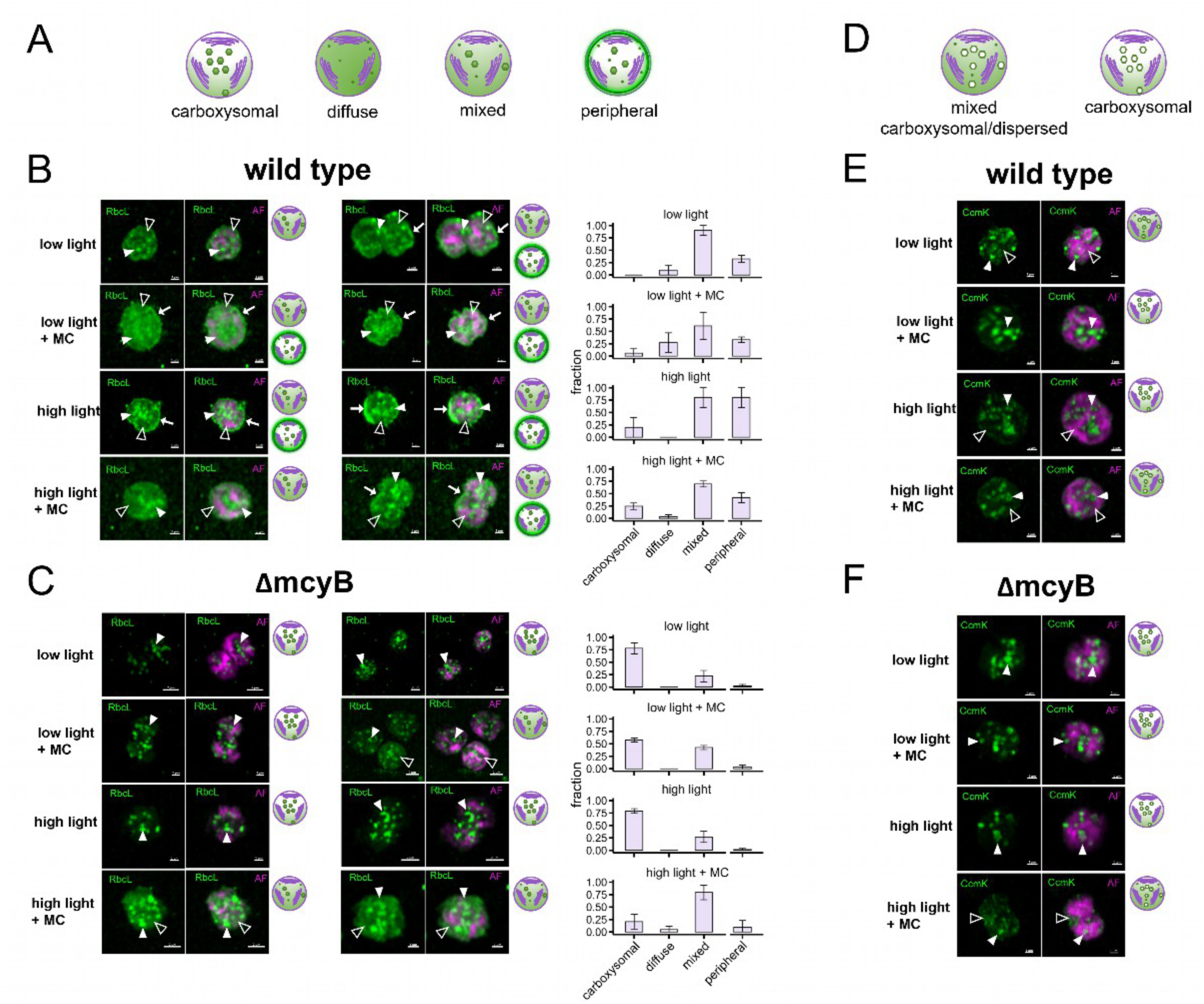
Intracellular localization of RubisCO and carboxysomes in *Microcystis* following microcystin addition. The RubisCO large subunit (RbcL) and the major carboxysome shell protein CcmK were detected using fluorescent antibodies in cells grown under low-light or high-light conditions, with (+MC) or without exogenous microcystin addition for 1h. (**A**) Schematic representation of the intracellular RubisCO distribution patterns identified by immunofluorescence microscopy. (**B**) Representative cells of the *Microcystis* wild type. (**C**) Representative cells of the MC-deficient mutant ΔmcyB. Cartoons (as in panel A) shown next to the micrographs illustrate the corresponding RbcL distribution patterns. Closed triangles point to distinct RbcL foci, presumed to represent carboxysomes, whereas open triangles denote diffuse cytoplasmic localization. Arrows highlight elongated stretches of peripheral RbcL (AF). Bar plots show the relative abundance of each distribution pattern. Quantification was based on 2–3 scanning regions per treatment, with 10–30 cells analyzed per region. (**D**) Schematic representation of carboxysome localization pattern identified by immunofluorescence microscopy. (**E**) Representative cells of the *Microcystis* wild type and (**F**) the MC-deficient mutant ΔmcyB. Cartoons (as in panel D) illustrate the corresponding CcmK localization patterns observed in each micrograph. Closed triangles indicate fully assembled carboxysomes, whereas open triangles denote diffuse cytoplasmic localization of CcmK.

A similar pattern was observed for carboxysome organization. In the wild type, the carboxysome shell protein CcmK frequently dispersed from discrete foci into more diffuse cytoplasmic regions, irrespective of light conditions (Fig. 4D). By contrast, carboxysome integrity was generally maintained in the MC-deficient mutant, with CcmK dispersal becoming apparent only after MC addition under high-light conditions (Fig. 4D). Together, these observations suggest that MC promotes a more dynamic organization of both RubisCO and carboxysome components, especially under high-light conditions.

## Discussion

In this study, we identify microcystin (MC) as a key contributor to the rapid acclimation of *Microcystis aeruginosa* PCC7806 to fluctuating irradiance. Compared with *Synechocystis*, *Microcystis* displayed a pronounced but transient response to high-light shifts, and this dynamic acclimation was strongly attenuated in the MC-deficient mutant. Because rapid adjustment to fluctuating light is thought to promote bloom formation and buoyancy regulation (*3*), MC likely contributes to the ecological success of *Microcystis*.

Although previous work already demonstrated enhanced high-light tolerance of the MC-producing wild type relative to the MC-deficient mutant (*12*), the underlying mechanism remained unresolved. In the present study, we therefore tested two hypotheses which are not necessarily mutually exclusive: first, that MC binding to RubisCO contributes mechanistically to enhanced high-light adaptation, and second, that extracellular MC functions as a signaling molecule governing the acclimation response. Although RubisCO activity showed pronounced light-dependent plasticity in *Microcystis*, our biochemical analyses provide little evidence for direct MC-mediated regulation. Instead, the lack of comparable regulation in the RMA strain suggests that other *Microcystis*-specific cellular factors modulate RubisCO activity, whereas MC likely acts upstream of carbon fixation rather than by directly altering RubisCO catalysis.

Consistent with a signaling role, extracellular MC induced a transient increase in 3PGA accumulation under high-light conditions *in vivo* and elicited a pronounced transcriptional response, particularly in the MC-deficient mutant. Transcriptomic analyses indicate that extracellular MC primarily targets the carbon acquisition machinery rather than RubisCO itself, thereby transiently enhancing inorganic carbon supply during high-light exposure. This response was substantially stronger than that observed in our previous study, where MC-dependent transcriptional effects were limited to a cryptic polyketide synthase biosynthetic gene cluster (BGC) following 24 h of low-light incubation (*10*). Together, these results demonstrate that MC signaling is highly dynamic and strongly dependent on environmental conditions.

Importantly, MC-induced regulation of carbon acquisition occurred as part of a broader transcriptional reprogramming. Although MC has frequently been linked to stress adaptation (*20*), it had limited effects on canonical oxidative stress markers. Instead, MC transiently and reciprocally regulated thioredoxin-related proteins, including a thioredoxin A variant and its associated CBS-CP12 protein implicated in CBB cycle regulation (*17*), while repressing the canonical *cp12*gene. These coordinated but opposing changes in functionally related proteins suggest a metabolic transition that may optimize CO_2_ fixation during high-light exposure. MC-dependent regulation also extended to photosynthetic gene expression, suggesting coordinated remodeling of carbon fixation and the photosynthetic apparatus. The transient nature of both the transcriptional response and photosynthetic stimulation supports the view that MC initiates a rapid acclimation program rather than establishing a stable physiological state (Fig. 5).

**Fig. 5:**
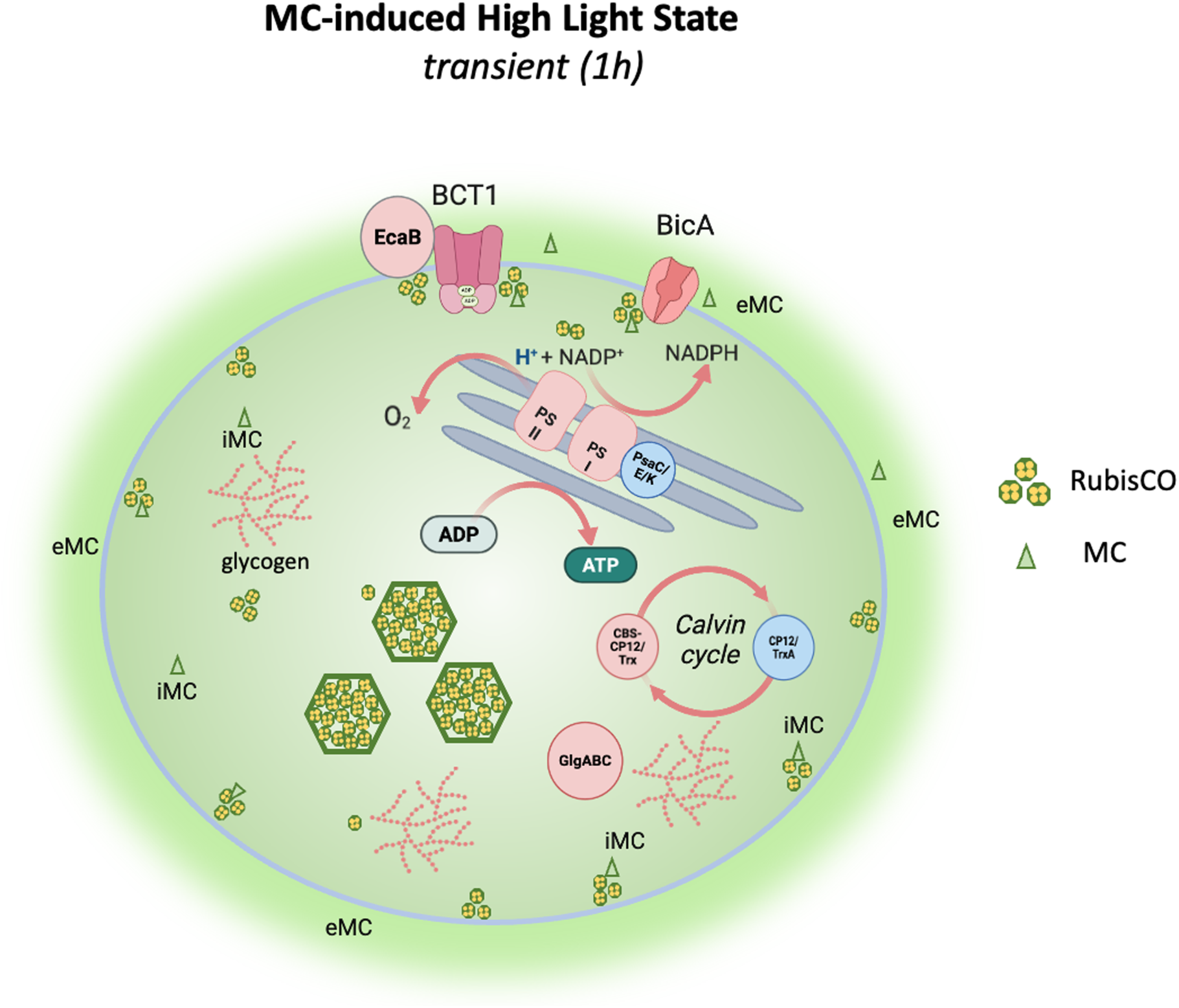
Model illustrating the transient MC-induced response of Microcystis one hour after transfer to high-light conditions. The illustration integrates data from physiological measurements, transcriptomic analyses, and *in situ* visualization. Cellular components shown in red and blue represent genes or processes that are transcriptionally upregulated or downregulated by MC, respectively.

Exogenous MC also partially restored the characteristic RubisCO and carboxysome organization of the wild type (*15*), demonstrating that extracellular MC regulates CCM architecture in addition to CCM gene expression. However, cytoplasmic membrane-associated RubisCO was not recovered, distinguishing extracellular signaling effects from intracellular MC-dependent processes (Fig. 5). Because this peripheral RubisCO state is closely associated with MC binding (*14*) it likely represents a distinct intracellular function of MC. Although the underlying mechanisms remain unclear, membrane-associated extracarboxysomal RubisCO in the wild type coincided with highly efficient glycogen accumulation. We therefore propose that this configuration, together with enhanced bicarbonate uptake, constitutes an alternative and potentially more flexible CCM that enables rapid responses to fluctuating environmental conditions. Such a system may contribute to the ecological success of *Microcystis* by facilitating transitions between periods of intense photosynthetic activity and subsequent glycogen utilization, processes that are crucial for buoyancy regulation of the colony-forming cyanobacteria. The signaling cascade underlying MC perception and extracellular release remains unknown, and future studies will be required to identify MC receptors or sensing mechanisms and determine whether global regulatory mechanisms mediate these responses. Notably, many MC-responsive genes encode hypothetical proteins (COG category R), highlighting the current limitations in assigning functions and emphasizing the need for further mechanistic studies.

Collectively, our findings identify MC as a coordinator of a rapid light-dependent CCM acclimation in *Microcystis*. The transient nature of MC-mediated responses may explain previously conflicting reports on its physiological functions and highlights the importance of considering dynamic regulatory processes when assessing the ecological role of MC, particularly under changing environmental and climatic conditions.

## Materials and methods

### Cultivation of cyanobacteria

Cyanobacterial liquid cultures were maintained in Erlenmeyer flasks in BG 11 medium (*21*) at 25°C, under constant white light illumination of 10 μE/m^2^s (i.e. “low light conditions”) at ambient CO_2_. The mutant strains *Microcystis aeruginosa* PCC7806 Δ*mcyB* and *Synechocystis sp.* PCC6803 RMA were cultivated with chloramphenicol at a final concentration of 5 µg/µl. Axenicity of cyanobacterial cultures was regularly assessed by streaking aliquots on solid R2A agar medium and subsequent 24h incubation at 30°C, 37°C and 22°C. For high light experiments, cultures were grown at constant white light illumination of 250 μE/m^2^s unless stated otherwise. Growth curves were recorded from liquid cultures grown in biological triplicates in 100 mL Erlenmeyer flasks under different light conditions. Samples of 1 mL were taken under sterile conditions and optical density at 750 nm (OD_750_) was measured photometrically.

### Mutagenesis of cyanobacteria

The microcystin deficient Δ*mcyB* mutant of *Microcystis aeruginosa* PCC 7806 was generated previously (*22*). The RMA (“<u>R</u>ubisco from <u>M</u>icrocystis <u>a</u>eruginosa”) mutant of *Synechocystis sp.* PCC6803 that has its native *rbcLXS* operon replaced by the corresponding sequence from *Microcystis aeruginosa PCC7806* was generated through natural transformation of an engineered plasmid vector harboring a template for gene replacement by homologous recombination. This plasmid was constructed as follows: All PCR reactions were carried out with Q5 polymerase (New England Biolabs), primer sequences are in table S1. All PCR amplificates were excised and purified from agarose gels with the GeneJet Gel Extraction Kit (ThermoFischer Scientific). DNA constructs used for mutagenesis were generated by Gibson assembly with the NEBuilder HiFi kit (New England Biolabs). The *rbcLXS* region of *Microcystis aeruginosa* PCC7806 (IPF_2532-IPF_2531-IPF_2530) was PCR amplified from genomic DNA with primers A and B. The upstream and downstream homology regions were PCR amplified from genomic DNA isolated from *Synechocystis sp.* PCC6803 with the primers C, D, E and F. A chloramphenicol resistance cassette was PCR amplified from the cloning vector pACYC184 primers G and H. The fragments were assembled into a pUC/pBR32 derived cloning vector, successful construction of the mutagenesis plasmid was verified by Sanger sequencing. Natural transformation of *Synechocystis sp.* PCC6803 was carried out by established protocols (*23*) and verified by PCR (Figure S1).

### Microcystin addition experiments

Commercially available Microcystin-LR (Enzo life sciences, product no. ALX-350-012-C100) was added to liquid cultures of *M. aeruginosa* grown in a Multi-Cultivator MC 1000-OD system (Photon Systems Instruments, Czech Republic). Samples were collected for quantification of 3-Phosphoglycerate and 2-phosphoglycolate (7 mL), RNA isolation for transcriptome analysis (2 mL), immunofluorescence microscopy (2 mL). Of each strain, three biological replicates were inoculated in fresh liquid medium at an OD_750_ of 0.1 and pre-cultivated in Erlenmeyer flasks under standard maintenance conditions to an OD_750_ of 0.4.

From each triplicate, two 80 mL aliquots were each transferred to one MC 1000-OD cultivation tube. Triplicates of each strain were pooled in equal parts yield an 80 mL pooled low light control sample. In total, eight MC 1000-OD tubes were prepared per strain: four for MC-LR addition (three triplicates and one low-light control) and four corresponding solvent controls. Liquid cultures were then pre-cultivated in the MC 1000-OD for adaptation for 24 h, 25°C, under low-light with gentle air bubbling. The experiment was started by increasing illumination to high light (250 μE/m^2^s), low light controls remained at 10 μE/m^2^s.

Concomitantly, MC-LR was added to a final concentration of 100 ng/mL, a corresponding volume of methanol was added to the solvent control samples. Samples were taken at the start of the experiment and after 1 h, 2 h and 3 h of incubation. Samples were centrifuged for 15 minutes at 4000 g at RT.

### Quantification of 3-phosphoglycerate by LC-MS

Cell pellets and supernatants for LC-MS quantification were freeze dried and processed as described (*24*). Compound quantification was performed after running extracts over a pentafluorophenylpropyl column (Supelco Discovery HS FS, 3 μm, 150 × 2.1 mm) with a mobile phase containing 0.1% formic acid on an LC–MS-8050 system (Shimadzu, Kyoto, Japan) and the incorporated LC–MS/MS method package for primary metabolites (version 2, Shimadzu). Compounds were eluted at 0.25 ml min^−1^ with a gradient of 1 min 0.1% formic acid, 95% aqua dest, 5% acetonitrile, within 15 min linear gradient to 0.1% formic acid, 5% aqua dest, 95% acetonitrile, 10 min 0.1% formic acid, 5% aqua dest, 95% acetonitrile with continuous injection electrospray ionization. Multiple reaction monitoring values given in the method package and the LabSolutions software (Shimadzu) sere used for compound identification and quantification. Authentic standards (Sigma-Aldrich) were included in all batches and used for calibration.

### RNA isolation and transcriptome analysis

For total RNA isolation, biological triplicates of each sampling event were pooled and RNA was isolated by the TRIzol (Thermo Fisher) method, precipitated with 300 mM Na-acetate (pH 5.2) and excess isopropanol, dissolved in water and stored at –80°C until further use. Sequencing of rRNA depleted total RNA was conducted by Biomarker Technologies (Münster, Germany) on an Illumina NovaSeq X plus platform by NovaSeqTM X Series25B Rgt Kit and paired-end 150 bp reads were generated. Reads were mapped to the genome sequence of *Microcystis aeruginosa* PCC7806 (accession CP020771.1). Differential gene expression analysis was carried out with edgeR. A list of software packages and parameters used is given in table S2.

### Quantification of intracellular glycogen content

For the quantification of total intracellular glycogen, 50 mL of biological triplicate liquid cultures of an OD_750_ of 0.5 were pre-incubated in Erlenmeyer flasks at low light conditions for 14 days to deplete cells of endogenous glycogen. Samples of 1 mL were taken before and after cultures were shifted to high light conditions. immediately pelleted and snap-frozen in liquid nitrogen. Cell pellets were extracted and glycogen quantified with a commercial kit (Glycogen Assay Kit, MAK016-1KT, Sigma-Aldrich).

### Quantification of oxygen evolution

Oxygen evolution of liquid cultures before and after onset of high light illumination was measured in biological triplicate cultures of an OD_750_ of 0.5 incubated for 14 days under low light conditions in Erlenmeyer flasks. For the measurement, culture aliquots were transferred to sterilized 2 mL glass HPLC vials. A needle-type oxygen sensor (preSense NTH-PST7 with a Microx 4 hand device) was inserted through a septum placed in the vial lid and oxygen concentration was recorded for 3 seconds.

### Immunofluorescence microscopy

Immunofluorescence microscopy was carried out as described previously (*15*) with the following modifications: Cell pellets were washed three times with PBS (137 mM NaCl, 2.7 mM KCl, 10 mM NaH_2_PO4, 1.8 mM KH_2_PO4, pH 8.0) and fixed in 4% para-formaldehyde/PBS (Thermo-Fisher) for 30 minutes at room temperature. Microscopy glass slides were sonicated for 20 minutes in ethanol and air dried, fixed samples were washed three times with PBS and resuspended in 200 µL water, 20 µL were spread on prepared glass slides, air dried and treated with 2 mg/mL lysozyme. Slides were blocked with 1% PVP K-30 in PBS-T (PBS with 0.3% Tween-20), washed and primary antibodies were applied at 4°C over night in a humid chamber. Primary antibodies used were: anti-Rubisco large subunit form I (anti-RbcL raised in chicken, dilution: 1:300, AS01 017 Agrisera, Vännas, Sweden) and anti-CcmK2, (custom rabbit polyclonal sera against the *Microcystis* protein, used at 1:200, for details see (*24*). After washing, samples were hybridized with the following secondary antibodies for 24 h at 4°C in the dark: Alexa Fluor 488 goat anti-rabbit (1:200) and Alexa Fluor 546 goat anti-chicken (1:200) (both Thermo Fisher Scientific). Samples were washed and air-dried, mounted in ProLongTM Glass Antifade Mountant (Thermo Scientific) and stored at –20°C until imaging. Images were captured with a Zeiss LSM 780 confocal laser scanning microscope. Alexa Fluor 488 was excited at 488 nm (detection spectrum 493–556 nm), Alexa Fluor 546 at 561 nm (570–632 nm) and autofluorescence at 633 nm (647–721 nm). The ZEN software was used for image analysis. For quantification of RbcL distribution patterns, 2-3 spatially separated regions were inspected per sample. Each region contained around 10-20 intact cells that showed unambiguous immunostaining. Spatial RbcL distribution was classified as either predominantly carboxysomal or predominantly diffuse. If both patterns were found, the cell was classified as mixed. Additionally, cells were classified according to the occurrence of continuous stretches (> 0.5µm) of peripheral RbcL localization.

### RubisCO purification from cell extracts

RuBisCO was isolated from 350 mL cyanobacterial liquid cultures grown in 1 L Erlenmeyer flasks under maintenance conditions (for low light samples) or at 150 μE/m^2^s (for high light samples) as previously described (*24*). Briefly, cell pellets were washed with water, resuspended in 20 mL RuBisCO extraction buffer (10 mM Bicine, 1 mM EDTA, 1 mM DTT, pH 8.1). Cells were lysed with a Constant cell disruptor (T-series; Constant Systems, UK) and debris was pelleted. The supernatant was subjected to two rounds of a fractionated ammonium sulfate precipitation, first at 20% saturation, the resulting supernatant then at 50% saturation. The resulting Rubisco-enriched pellet was resuspended in FPLC buffer A (100 mM K_2_HPO_4_, 1 mM EDTA, 1 mM DTT, pH 7.6), treated with 0.5% w/v Triton-X, precipitated with 20% w/v PEG6000 and subjected to anion-exchange FPLC with a MonoQ 4.6/100 PE column on an Äkta pure system (both Cytiva) with a linear gradient of 1 M KCl in buffer A, run for 50 column volumes from 0 mM to 500 mM KCl. RubisCO typically eluted at a concentration of 250-350 mM KCl. RuBisCO fractions were stored for imminent use in 50% saturated ammonium sulfate solution at 4°C or snap frozen in liquid N_2_ and stored at –80°C. Purity was assessed by SDS-PAGE and RbcL and RbcS immunoblots, concentration was determined by a Bradford assay (Thermo Fisher)

### RuBisCO activity assay in vitro

RubisCO carboxylase activity was measured essentially as described previously (*24*). RubisCO concentration weas adjusted to 0.2 µg/µl in carboxylation buffer (20 mM Bicine, 50 mM MgCl_2_, 50 mM NaHCO_3_, pH 8.0) and activated for 10 minutes at 25°C. Carboxylation was initiated by adding Ribulose-1,5– Bisphosphate (tetra-sodium hydrate, >99% purity, Sigma-Aldrich, 83895) to a final concentration of 0.4 mM.. Samples were collected after 30 s, 60 s, 120 s, 240 s and immediately transferred to 10 M formic acid to stop the reaction. Samples were vacuum dried and 3PGA was quantified by LC-MS. To assess the influence of MC-LR on RubisCO activity, RubisCO isolates were incubated in the presence of 0.5 mM TCEP (pH9.0) and 2 ng/µL MC-LR for 24 h at 4°C prior to activity assays. Control reactions had the solvent (methanol) added instead of MC-LR.

### Mass photometry of RubisCO isolates

Microscope coverslips (1.5 H, 24 x 60 mm, Carl Roth) and CultureWellTM Reusable Gaskets (CW– 50R-1.0, 50-3mm diameter x 1 mm depth) were cleaned with three consecutive rinsing steps of ddH2O and isopropanol, then dried under a stream of pressurized air. For measurements, gaskets were assembled on coverslips and placed on the stage of a OneMP mass photometer (MP, Refeyn Ltd, Oxford, UK) with immersion oil. Assembled coverslips were held in place using magnets. For measurements, gasket wells were filled with 18 μL 1x phosphate-buffered saline (PBS, 10 mM Na2HPO4, 1.8 mM KH2PO4, 137 mM NaCl, 2.7 mM KCl, pH 7.4) to enable focusing of the glass surface. After focusing, 2 μL sample were added, rapidly mixed while keeping the focus position stable and measurements were started (<15 s after sample addition). Data was acquired for 60 s at 100 frames per second using AcquireMP (Refeyn Ltd, v1.2.1). Samples were prepared by diluting purified protein to 10 μM monomer concentration in desalting buffer (25 mM Tricine (pH 8.0), 75 mM NaCl), as determined by absorption at 280. Immediately prior to measuring, samples were further diluted to a final concentration of 500-1000 nM and 2 μL thereof were mixed with 18 μL PBS that had been used to focus the MP on the glass slide. MP contrast values were calibrated to molecular masses using commercial NativeMarkTM Unstained Protein Standard (Thermo Fisher) as a standard. For calibration, 1x NativeMark was diluted 30– fold in 1x PBS and subsequently used as a 10x stock (2 μL standard added to 18 μL 1x PBS).

Calibration measurements showed well resolved peaks at contrast corresponding to 66 kDa, 146 kDa, 480 kDa and 1048 kDa, which were integrated and fit to a linear regression using DiscoverMP (Refeyn Ltd, v1.2.3). All MP images were processed and analyzed using DiscoverMP (Refeyn Ltd, v.1.2.3).

### Determination of specificity coefficient of RubisCO isolates

CO_2_/O_2_ specificity assays were performed as published previously (*25*). In short, purified RubisCOs were incubated in 20 mL septum capped glass scintillation vials containing 1 mL 30 mM triethanolamine (pH 8.3), 15 mM magnesium acetate, and 0.01 mg/mL carbonic anhydrase. Reaction mixtures were pre-equilibrated in defined gas mixtures containing either 995.009 ppm O_2_ and 4991 ppm CO_2_ (for low specificity variants) or 999.300 ppm O_2_ and 700 ppm CO_2_ (for high specificity variants, Air Liquide, Singapore) before initiating the reactions by the addition of [1-3H]-RuBP. After running to completion, products were dephosphorylated using alkaline phosphatase (10 U/reaction), separated on an HPX-87H column (Bio-Rad) and peaks were quantified using liquid scintillation counting. The SC/O was subsequently calculated as described previously (*26*).

## Supporting information

SI Section

## Acknowledgments

We are grateful to Lena Uhlig (University of Potsdam), Klaudia Radloff-Michel and Manja Henneberg (University of Rostock) for technical assistance. We further thank Cheryl Kerfeld for critical discussion of the manuscript.

## Funding

DFG project number 239748522– SFB 1127 (ED)

DFG project number 467315902 (ED)

DFG project number INST 264/125-1 FUGG (MH)

## Author contributions

Examples: Conceptualization: ED, AG

Methodology: ED, AG, GH, MH

Investigation: AG, VW, LS, ST

Visualization: ED, AG, VW, LS

Funding acquisition: ED, GH, MH

Writing – original draft: ED, AG

Writing – review & editing: ED, AG, VW, LS, ST, GH, MH

## Competing interests

Authors declare that they have no competing interests.

## Data, code, and materials availability

RNASeq transcriptomic data is available as trimmed, raw reads in the public NCBI database under the bioproject accession number PRJNA1490535. All raw data for microscopy, LC-MS, mass photometry, kinetic measurements as well as plasmid vectors and cyanobacterial mutant strains are available from the corresponding author upon request. All other data are available in the manuscript or the supplementary materials.

## Supplementary Materials

Figs. S1 to S5

Tables S1 to S2

Data S1

## Notes

https://www.ncbi.nlm.nih.gov/sra/PRJNA1490535

### Competing Interest Statement

The authors have declared no competing interest.

