## Supplementary material for "Microcystin-Driven Control of the Carbon-Concentrating Mechanism Shapes CO_2_ Fixation Dynamics in *Microcystis aeruginosa* PCC 7806": SI Section

### Content

### 1. Supplementary tables

**Table S1. Primers used in this study.**
**Table S2. Software and parameters used in transcriptome analysis.**

### 2. Supplementary figures

**Figure S1. Confirmation PCR(S) RMA mutant.**
**Figure S2. RubisCO-MC binding blot, without and with MC addition**
**Figure S3. Global transcriptional response to external microcystin addition differentiated by COG gene ontology.**
**Figure S4. Transcriptional response of selected genes to external microcystin addition.**
**Figure S5. IFM controls**

### 3. Dataset S1

**S1 comprehensive RNASeq data file.xlsx**

### 1. Supplementary Tables

**Table S1**. Primers used in this study for Gibson Assembly.

| Name | Sequence (5' to 3') | Description |
|---|---|---|
| A | ATGGTGCAAGCCAAATCCAA | amplification of *rbcLXS* from *Microcystis aeruginosa* PCC7806 |
| B | TTAGTAGCGGCCAGCATTG | amplification of *rbcLXS* from *Microcystis aeruginosa* PCC7806 |
| C | TGGGGTGGATGCAGTGGGCCCAAGCAGGTCAATCGGTGGGTTACA | amplification of *rbcLXS* upstream homology region from *Synechocystis* sp. PCC6803 |
| D | AACCTTTGGATTTGGCTTGCACCATCTAGGTCAGTCCTCCATAAA | amplification of *rbcLXS* upstream homology region from *Synechocystis* sp. PCC6803 |
| E | ATTTTTTTAAGGCAGTTATTGGTGCCCGTTACAGTTTTGGCAATTAC | amplification of *rbcLXS* downstream homology region from *Synechocystis* sp. PCC6803 |
| F | TCAAACACTGATAGTTTAAACAGGTTGAGGGTCAGGAGCATATCC | amplification of *rbcLXS* downstream homology region from *Synechocystis* sp. PCC6803 |
| G | TAAACCCAATGCTGGCCGCTACTAAAAGAGGTTCCAACTTTCACCA | amplification of *CmR* resistance cassette from pACYC184 |
| H | GGGCACCAATAACTGCCTTA | amplification of *CmR* resistance cassette from pACYC184 |

**Table S2**. Software and parameters used in transcriptome analysis.

| Software | Version | Main parameter |
|---|---|---|
| bowtie2 | 02.02.2004 | -p 6 --very-sensitive |
| RSEM | 01.02.2019 | --bowtie2 --paired-end -p 12 |
| diamond | 2.0.15 | -k 100 -e -evalue 1e-5 -f 5 |
| InterProScan | 5.34-73.0 | -appl Pfam -goterms -iprlookup -pa -f xml -dp -t p |
| hmmscan | 03.03.2002 | --noali --cut_nc --acc --notextw |
| edgeR | 3.32.1 | dispersion=0.01 |
| R/clusterProfiler | 04.04.2004 | minGSSize=1,maxGSSize=10000,pAdjustMethod="fdr" |
| R/topGO | 2.48.0 | firstSigNodes=5 |

### 2. Supplementary Figures

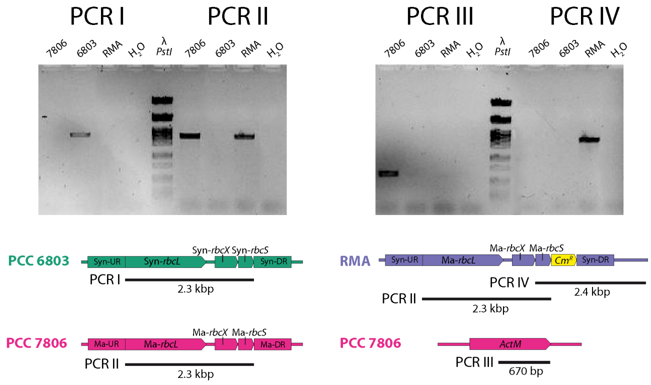


**Figure S1: Confirmation PCR(S) RMA mutant.**

The *rbcLXS* locus of 3 cyanobacterial strains was genotyped by PCR with isolated genomic DNA using specific primers. The RMA mutant of *Synechocystis* carries a homozygous replacement of *rbcLXS* from *Microcystis aeruginosa* PCC7806 at the native locus.

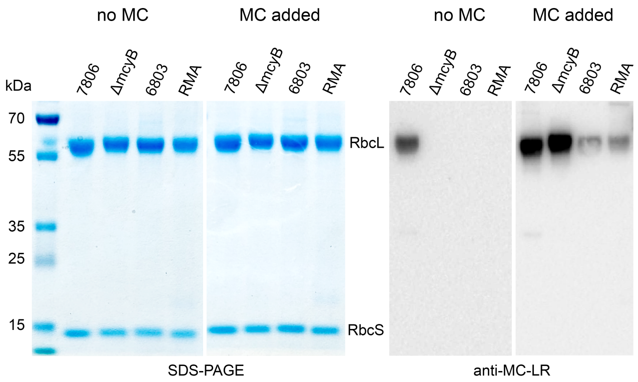


**Figure S2: RubisCO-MC binding blot, with and without MC addition**

Purified RubisCO from four different cyanobacterial strains was subjected to SDS-PAGE (left) and anti-microcystin immunoblotting (right) before and after MC addition. These RubisCO were then used for activity measurements shown in figure 2. MC-bound RubisCO was active but prone to aggregation and largely resistant to protease digestion. Therefore, the MC binding site on the RubisCO complex could not be determined experimentally. Strain codes are: “7806” for the *Microcystis aeruginosa* wild type, “ΔmcyB” for its microcystin-free mutant, “6803” for the *Synechocystis* sp. wild type and “RMA” for a *Synechocystis* RubisCO replacement mutant that expresses the *Microcystis* variant of *rbcLXS*.

| COG class legend | |
|---|---|
| X | Mobilome: prophages, transposons |
| V | Defense mechanisms |
| R | General function prediction only |
| C | Energy production and conversion |
| J | Translation, ribosomal structure and biogenesis |
| M | Cell wall/membrane/envelope biogenesis |
| O | Posttranslational modification, protein turnover, chaperones |
| T | Signal transduction mechanisms |
| H | Coenzyme transport and metabolism |
| E | Amino acid transport and metabolism |
| P | Inorganic ion transport and metabolism |
| S | Function unknown |
| L | Replication, recombination and repair |
| Q | Secondary metabolites biosynthesis, transport and catabolism |
| K | Transcription |
| G | Carbohydrate transport and metabolism |
| I | Lipid transport and metabolism |
| F | Nucleotide transport and metabolism |
| D | Cell cycle control, cell division, chromosome partitioning |
| U | Intracellular trafficking, secretion, and vesicular transport |
| N | Cell motility |
| W | Extracellular structures |
| Z | Cytoskeleton |

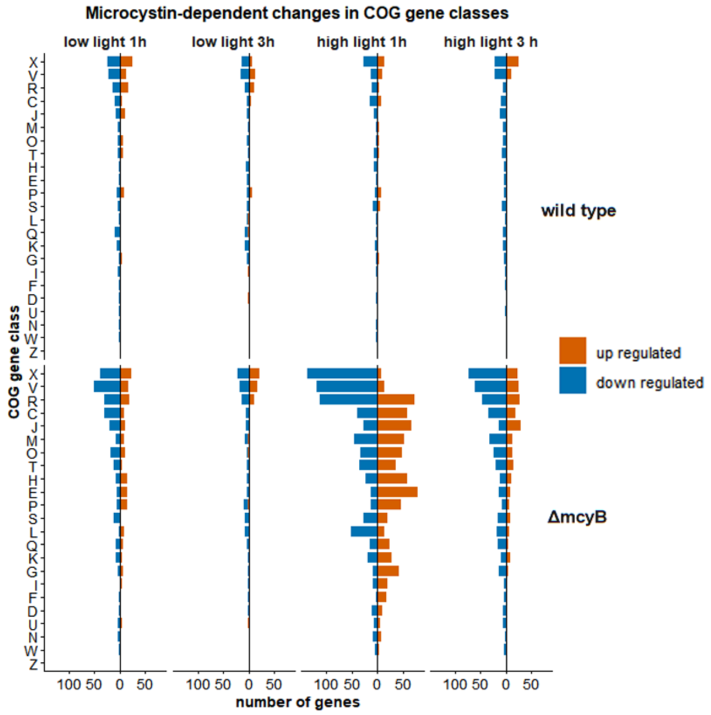


**Figure S3: Global transcriptional response to external microcystin addition differentiated by COG gene ontology.**
Number of genes responding to microcystin addition to liquid cultures of the *Microcystis* wild type and its microcystin-free mutant ("ΔmcyB") grown under low light or high light for 1 hours and 3 hours. Orange bars facing right show up regulated genes, blue bars (facing left) show down regulated genes with the most responsive gene classes sorted to the top.

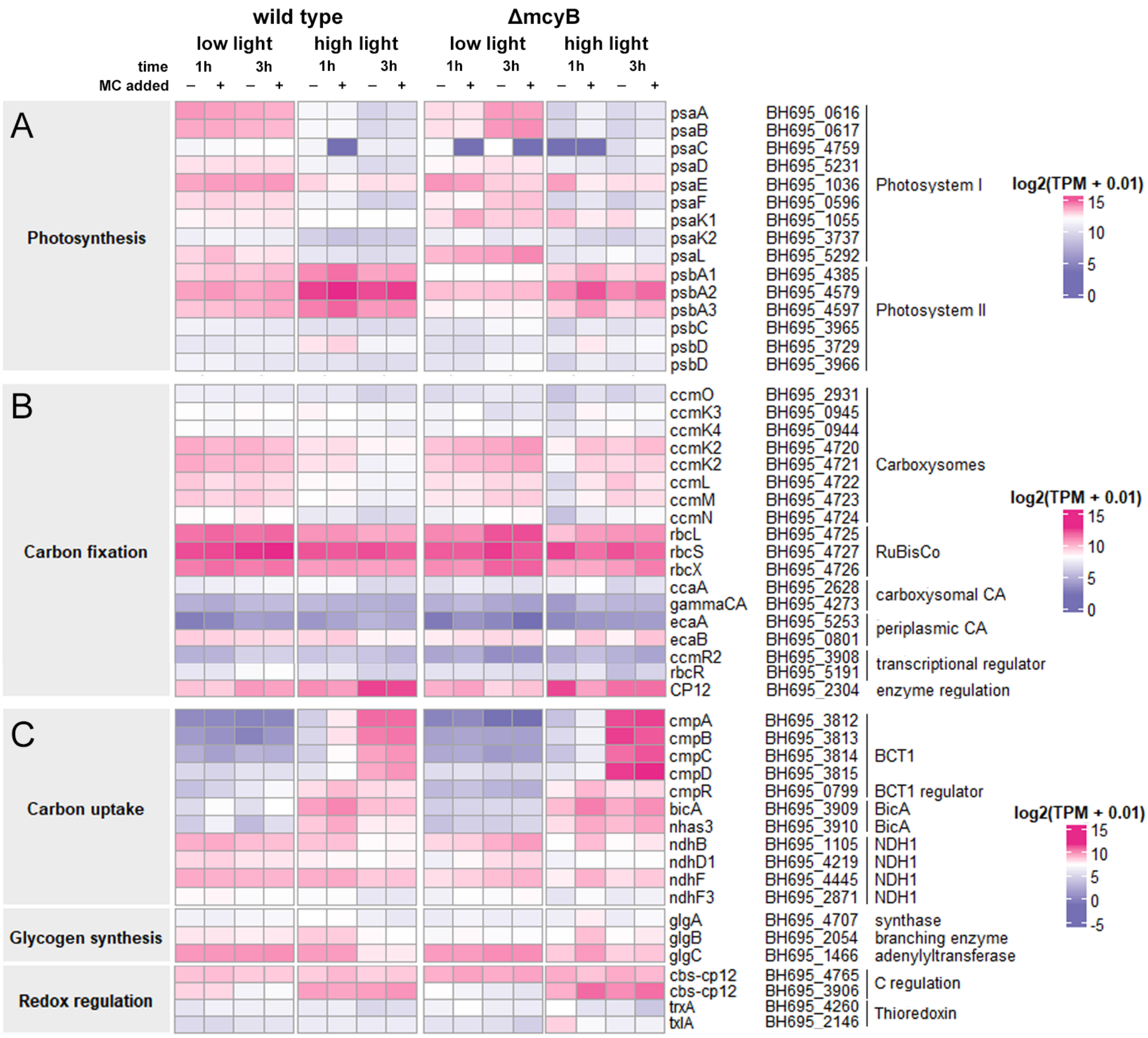


**Figure S4: Transcriptional response of selected genes to external microcystin addition.**
Heat map of selected genes displaying $log_2$-transformed transcript levels in TPM (transcripts per million) from cells grown with ("+") and without ("-") externally supplied microcystin (control group). Biological replicate strains of *Microcystis aeruginosa* PCC7806 wild type and its microcystin-free mutant ("ΔmcyB") were grown under low light and high light for 1 hours and 3 hours. Broad biological gene classification is given in shaded boxes (left), gene names, locus tags and functional gene subgroups (in that order) are shown to the right. A: genes related to photosynthesis, B: genes related to carbon fixation, C: genes related carbon uptake, glycogen synthesis and redox regulation.

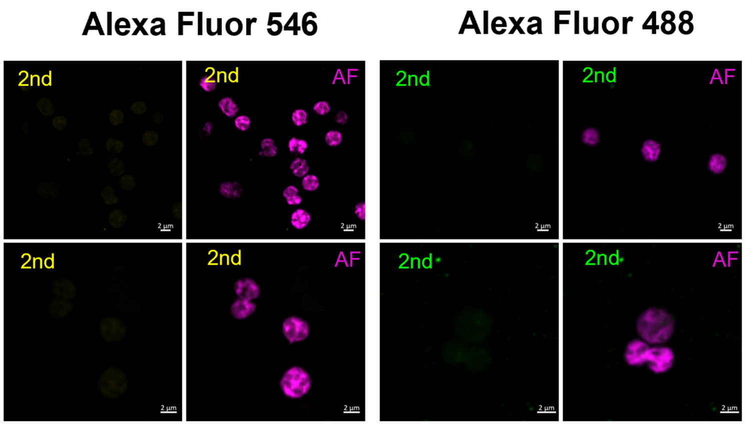


**Figure S5: IFM controls.**
Control hybridizations of *Microcystis* cells and fluorescent secondary antibodies ("2nd") used for immunofluorescence microscopy in this study.
